# Cross-view graph neural networks for spatial domain identification by integrating gene expression, spatial locations with histological images

**DOI:** 10.1101/2024.07.25.605067

**Authors:** Songyan Liu, Yin Guo, Zixuan Zhang, Shuqin Zhang, Limin Li

## Abstract

The latest developments in spatial transcriptomics technology provide an unprecedented opportunity for in situ elucidation of tissue structure and function. Spatial transcriptomics can provide simultaneous, multi-modal, and complementary information, including gene expression profiles, spatial positions, and histological images. Despite these capabilities, current methodologies often fall short in fully integrating these multi-modal datasets, thereby limiting their ability to fully understand tissue heterogeneity. In this study, we propose XVGAE (cross-view graph autoencoders), a novel approach that integrates gene expression data, spatial coordinates, and histological images to identify spatial domains. XVGAE constructs two distinct graphs: a spatial graph from spatial coordinates and a histological graph from histological images, and these graphs enable XVGAE to learn specific representations for each view and propagate information between them using cross-view graph convolutional networks. The experiments on benchmark datasets of the human dorsolateral prefrontal cortex show demonstrate that the XVGAE could achieve better clustering accuracy than state-of-the-art methods, and further experiments on four real spatial transcriptomics datasets on different sequencing platforms show that the XVGAE could identify biologically meaningful spatial domains with smoother boundary than other methods.

## 1 Introduction

Spatial transcriptomics (ST) is a recent breakthrough in biotechnogy [1], enabling simultaneous measurement of cell positions and comprehensive detection of cellular transcriptomes. It overcomes the limitations of single-cell sequencing by providing spatial context for cellular interactions [2]. ST technologies can be broadly classified into imaging-based (e.g., seqFISH [3], osmFISH [4], MERFISH [5]) and sequencing-based methods(e.g., 10x Visium [6], Slide-seq [7], Slide-seqV2 [8], Stereo-seq [9]). While imaging-based methods offer subcellular resolution, they may lack complete genome coverage. Conversely, sequencing-based approaches provide genome-wide expression profiling but typically capture spatial information for multiple cells within a single spot.

The development of clustering methods tailored for spatial transcriptomics data is currently at the forefront of biotechnology. It aids in identifying spatial domains, areas that exhibit spatial consistency in both gene expression and histology, forming the foundation for subsequent biological analyses [10]. Based on the utilization of spatial information, existing clustering methods for identifying spatial domains can be categorized into two types: non-spatial clustering methods and spatial clustering methods.

Traditional clustering methods, such as mclust, K-means [11], SC3 [12], and the Louvain [13] algorithm focus on gene expression data, neglecting spatial positional information and histological images. This omission often leads to discontinuities in their outcomes, which fail to align with the actual organizational structure. To leverage more information and enhance clustering results, novel algorithms have been developed to integrate spatial positional information and histological images, aiming to achieve more continuous and realistically aligned clustering outcomes. For instance, Giotto [14] identified spatial domains by employing the hidden Markov random field (HMRF) model to capture the spatial relationships between spots. BayesSpace [15], a fully Bayesian statistical approach, utilized information from spatial neighborhood information to enhance the resolution of spatial transcriptomics data for clustering analysis. STAGATE [16] combined gene expression and spatial information, detecting spatial domains through a graph attention autoencoder framework. SEDR [17] employed an autoencoder to construct a low-dimensional latent representation of gene expression, subsequently embedding the corresponding spatial information through a variational graph autoencoder. While these methods have improved clustering results to some extent, they overlook histological information.

Recently, some approaches have started incorporating histological image information to enhance spatial domain recognition. For example, stLearn [18] leveraged gene expression from neighboring points along with features from histological images to identify spatial domains. Similarly, SpaGCN [10] uses graph convolutional networks (GCNs) to recognize spatial regions by constructing a spatial graph that integrates both gene expression and histological information. While these methods can integrate histological image information, there are still limitations in accurately identifying spatial domains. This indicates that current approaches to integrating histological images may not be optimal. To fully leverage histological information, it can be beneficial to treat it as a distinct data modality and apply dedicated multi-view algorithms for integration.

To address this issue, we propose a cross-view graph autoencoder (XVGAE) framework for accurately identifying spatial domains by integrating spatial position, gene expression profiles, and histological images to learn low-dimensional latent embeddings. By constructing a graph from spatial locations and another graph from histological images, with the same node features of gene expression levels, the XVGAE can learn specific representations for each view and propagate information between them through our cross-view graph convolutional networks. We extensively demonstrate the effectiveness of this framework on spatial transcriptomics data from different platforms. Comparisons with existing algorithms indicates that XVGAE exhibits robust capabilities in accurately identifying spatial domains. Furthermore, it can effectively identify oncogenic genes, facilitating targeted analysis in cancer regions during downstream analysis. This is essential for understanding tumor heterogeneity and advancing personalized cancer treatment.

## 2 Results

In this section, we conducted experiments on four spatial transcriptomics datasets including human dorsolateral prefrontal cortex (DLPFC) dataset with manually labeled layers (10x Visium), mouse brain tissue (10x Visium), human breast cancer (10x Visium) and mouse olfactory bulb (Stereo-seq). We compared our XVGAE with other state-of-the-art methods, including STAGATE [16], BayesSpace [15], stLearn [18], and SEDR [17]), as well as SpaGCN [10], which is a widely used method that also integrates histological image information.

### 2.1 Overview of the XVGAE

The development of clustering methods tailored for spatial transcriptomics data is at the forefront of computational biology, aiding in identifying spatial domains that exhibit spatial consistency in both gene expression and histology. In this study, we introduce XVGAE (cross-view graph autoencoders), an innovative approach for identifying spatial domains by integrating gene expression data, spatial locations, and histological images. We construct two attributed graphs to capture the relationship among spots from two different views of spatial locations and histological images, with the corresponding adjacency matrices *A*_*P*_ and *A*_*I*_ being calculated from spatial locations *P* and preprocessed histological image features *X*_*I*_, respectively, and the same nodes with the same gene expression profiles as node attributes. The two attributed graphs, denoted as 𝒢_*P*_ = (*A*_*P*_, *X*) and 𝒢_*I*_ = (*A*_*I*_, *X*), represent the spatial and histological views of the spots.

XVGAE consists of a cross-view GCN encoder to capture hidden representations by integrating the two graphs, and a decoder to reconstruct the gene expression matrix *X*. The cross-view GCN encoder first employs two GCNs to extract view-specific features for the spatial graph 𝒢_*P*_ and the histological graph 𝒢_*I*_. Then, it integrates the view-specific information from both views by conducting cross GCNs on the adjacency matrix from one view with the view-specific representations from the other view. This mutual propagation of information between the spatial and histological views results in fused representations. In addition to the cross-view graph encoder, gene expression features are directly extracted from the gene expression matrix. The final representation is a combination of these gene expression features and the fused graph representations, with a learnable coefficient that is adjusted using gradient descent. The main idea of XVGAE method is illustrated in Fig. 1.

**Fig. 1.**
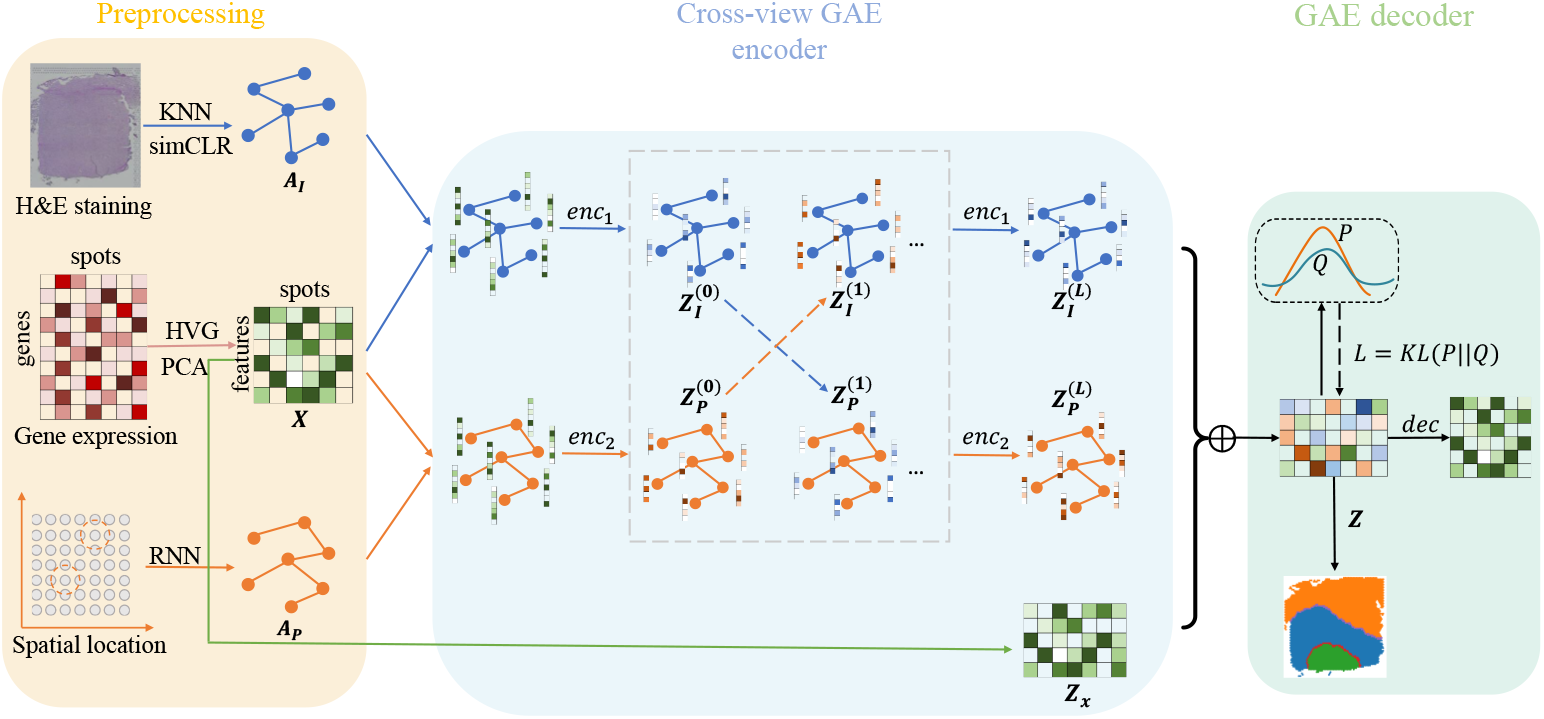
Overview of XVGAE: Given spatial transcriptomics data with three components: gene expression, spatial locations, and histological images as the input, the preprocessing step results in a spatial graph (*A*_*P*_, *X*) and a histological graph (*A*_*I*_, *X*), with nodes for both graphs are the same spots represented by gene expression matrix *X*. A cross-view GAE encoder and an AE encoder are simultaneously taken to extract latent features *Z* for spots, and a decoder is to reconstruct the gene expression matrix *X*. The loss function consists of a reconstruction loss of *X* and a clustering loss calculated from *Z*.

The effectiveness of XVGAE is extensively demonstrated on spatial transcriptomics data from different platforms. Comparisons with existing algorithms indicate that XVGAE exhibits robust capabilities in accurately identifying spatial domains and oncogenic genes, facilitating targeted analysis in cancer regions and advancing personalized cancer treatment.

### 2.2 XVGAE enhances the identification of established layers within the human dorsolateral prefrontal cortex dataset

We analyzed the 10x Visium dataset from the human dorsolateral prefrontal cortex (DLPFC) for spatial domain identification. This is a publicly accessible 10X Visium ST benchmark dataset, where Maynard et al [19] manually labeled the cortical layers (L1-L6) and white matter (WM) in 12 sections of the DLPFC data by genetic markers and cytoarchitecture (Fig. 2A for section DLPFC 151672 and Fig. S1 for the other 11 sections).

**Fig. 2.**
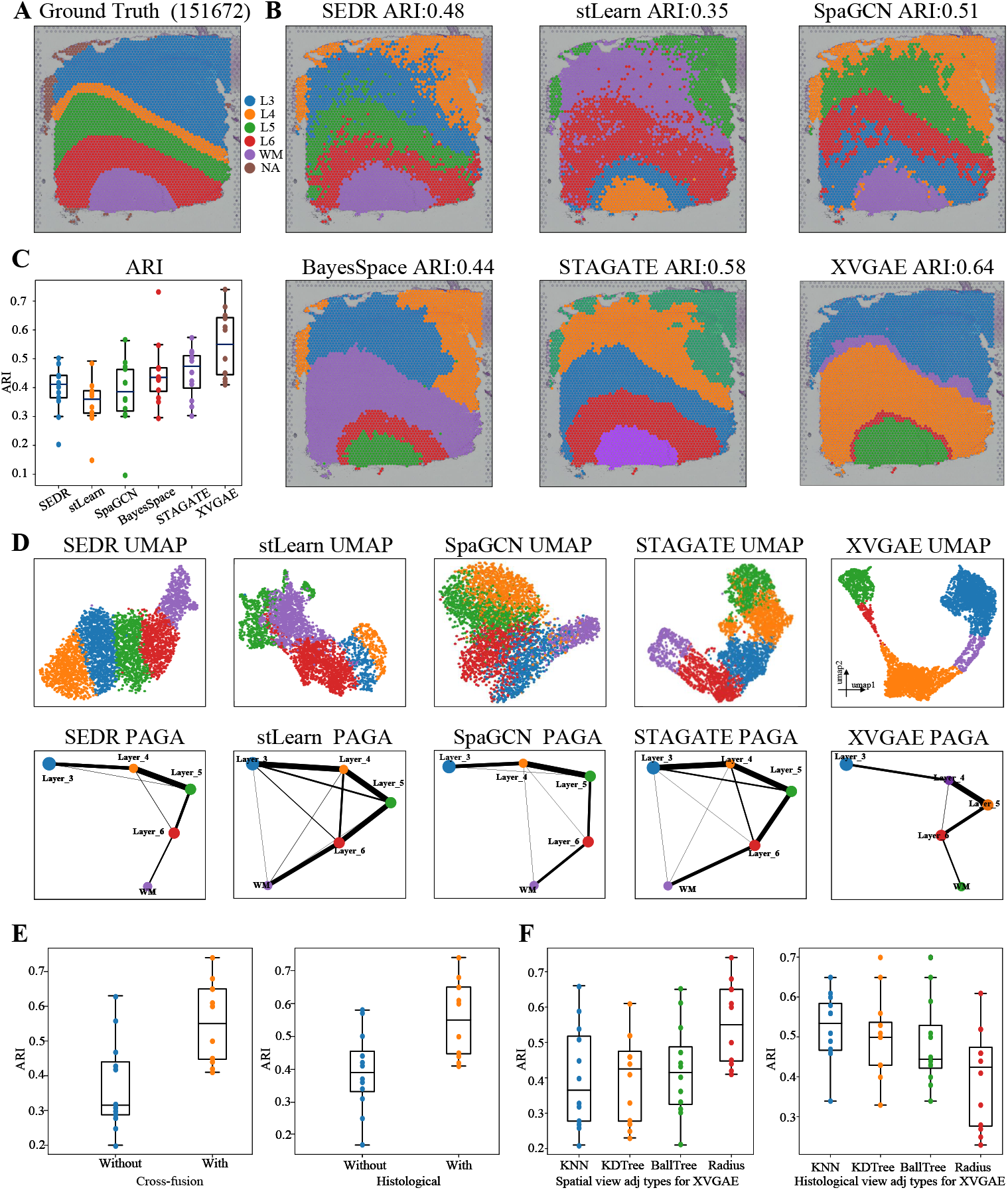
XVGAE enhances the identification of established layers within the human dorsolateral prefrontal cortex dataset. (A) Manual annotation of DLPFC section 151672. (B)Spatial domains identified by SEDR, stLearn, SpaGCN, BayesSpace, STAGATE and XVGAE on DLPFC section 151672. (C) ARI boxplot of the performance of XVGAE and other algorithms for all 12 DLPFC sections. Within the boxplot, the central line, box boundaries, and whiskers represent the median, upper and lower quartiles, and 1.5 times the interquartile range, respectively. (D) UMAP visualizations and PAGA graphs generated by SEDR, stLearn, SpaGCN, BayesSpace, STAGATE and XVGAE in the DLPFC section 151672. (E) ARI boxplots of whether using cross-view module and whether integrating histological information in XVGAE. (F) ARI boxplots of four methods for constructing adjacency matrices in XVGAE.

Fig. 2B illustrates the spatial domains identified by the XVGAE and other methods for section DLPFC 151672. The XVGAE shows superior performance in accurately delineating boundaries between different layers, achieving the highest Adjusted Rand Index (ARI) [20], whereas other methods exhibit varying degrees of blurring across different domains. For example, the SEDR, SpaGCN, and stLearn demonstrate significant noise, and all compared methods tend to confuse layer L3 with layer L4. Notably, only our XVGAE accurately identifies the very thin layer L4 in the annotation. Furthermore, the stLearn and BayesSpace incorrectly classify the WM layer as a combination of two distinct regions, whereas XVGAE accurately identifies the WM layer. Although SpaGCN [10] also integrates histological image information, its clustering boundaries tend to be discontinuous, with a significant number of outliers, thereby affecting the clustering accuracy.

Fig. 2C and Fig. S3 further report the ARI values and NMI values, respectively, obtained by different methods for all 12 sections in the form of box plots. The XVGAE achieves significant improvements on 12 sections compared to other recent methods. The median ARI for XVGAE is 0.55, notably higher than the second best method STAGATE, which achieves 0.47. The spatial domains identified by XVGAE are more consistent with the ground truth, indicating a substantial improvement over other methods, as illustrated in Fig. S1. This figure visualizes the spatial domains for all the other 11 sections by different methods, where XVGAE consistently achieves the highest ARI scores across multiple sections. For sections 151669-151672, which share similar morphological characteristics, XVGAE performs exceptionally well. While other methods fail to identify the complete layer L1 in these sections, XVGAE accurately distinguishes layer L1 from other regions. For instance, in DLPFC section 151669, XVGAE clearly outlines layer boundaries and achieves the highest clustering accuracy (ARI=0.71). These experimental results highlight XVGAE’s superiority in spatial domain recognition and effective integration of image information.

Figure 2D presents the UMAP and PAGA [21] trajectory plots derived from embeddings generated by XVGAE and other methods for section 151672, and the corresponding plots for the other 11 sections are reported in Fig. S2. The UMAP visualizations show that SEDR and XVGAE successfully distinguish between different categories, highlighting the functional similarities among adjacent cortical regions. In the PAGA trajectory plots, the edges connecting nodes signify potential transitions or relationships between clusters, with thicker edges indicating stronger evidence for these inter-cluster transformations. By analyzing these thicker edges, one can infer the developmental trajectories of spots across various spatial domains. Notably, only the PAGA trajectory plot generated by XVGAE aligns with the order of the spatial locations of the identified spatial domains, clearly illustrating a coherent developmental trajectory. This demonstrates that the embeddings learned by XVGAE retain biologically meaningful information, providing a more accurate and insightful representation of spatial transcriptomics data.

We further conducted three ablation studies on the DLPFC dataset and calculated their ARI values to evaluate the effectiveness of different modules in XVGAE. These studies particularly highlight the importance of the cross-view module and the inclusion of histological image data. Treating histological images as a separate view for learning and effectively integrating them with information from other modalities is a prominent innovation of XVGAE, which significantly impacts its performance. To investigate the influence of histological images on the XVGAE’s performance, we ran XVGAE separately with and without histological images. Specifically, when not using histological images, only the gene expression matrix and spatial location information are provided as inputs. In this scenario, the adjacency matrices for both views are constructed based on spatial proximity relationships, and node features were derived from the gene expression matrix, resulting in two identical views. In Fig. 2E, we presented the clustering results separately. Experimental results demonstrate that the utilization of histological image information effectively improves the clustering accuracy of XVGAE.

Next, we conducted experiments with or without the cross-view GNN module to explore its contribution on the clustering accuracy of XVGAE. Specifically, For the case without the cross-view GNN module, we simply employed only two layers of GCN to learn from two different views, while our designed cross-view module takes the latent features from the previous layer of the other view as input when calculating the latent features for the next layer. Fig. 2E shows that the cross-view GNN module significantly enhances the clustering performance of XVGAE. This demonstrates the role of the cross-view GNN module in ensuring the effective integration of information from different views, making it a key component in leveraging histological image information and improving clustering accuracy.

Finally, we evaluated the impact of different methods for constructing adjacency matrices on model accuracy. We designed experiments using four methods [22](KNN, KDTree, BallTree, Radius) to construct adjacency matrices for the two views and calculated their ARI values, as shown in Fig. 2F. Notice that different adjacency matrix construction methods would affect the performance of XVGAE. When constructing the adjacency matrix based on spatial proximity relationships, employing Radius yielded the best clustering performance. Conversely, using KNN to construct the adjacency matrix based on histological image similarity resulted in higher ARI and better model robustness.

### 2.3 XVGAE identifies better spatial domains for mouse brain

We evaluated XVGAE on a more complex spatial transcriptomics dataset of mouse brain based on 10x Visium technology. This dataset differs from DLPFC in that its spatial domains are more finely delineated. It contains two slices: the Sagittal-Posterior slice (MBP) and the Coronal slice (MBC). The corresponding anatomical reference is annotated by the Allen Mouse Brain Atlas [23]. The histological images and the Allen Mouse Brain Atlas are shown in Fig. 3A. In the experiment, since the ground truth categories were unknown, we clustered the embeddings from various methods separately for mouse brain sagittal and coronal slices into multiple distinct categories. We presented the clustering results for different methods using varying numbers of clusters (11, 15, 17, 20, 22, 24) in Fig. S6 and S7 for mouse brain sagittal-posterior and coronal slices, respectively. Additionally, we highlighted the selected clustering results of different methods that exhibited clearly identified regions in Fig. 3B-C for both slices, where the cluster numbers for SEDR, stLearn, SpaGCN, STAGATE, and XVGAE are 16, 16, 11, 11, and 18 for the MBP data shown in Fig. 3B, and 20, 16, 18, 24, and 25 for the MBC data shown in Fig. 3C.

**Fig. 3.**
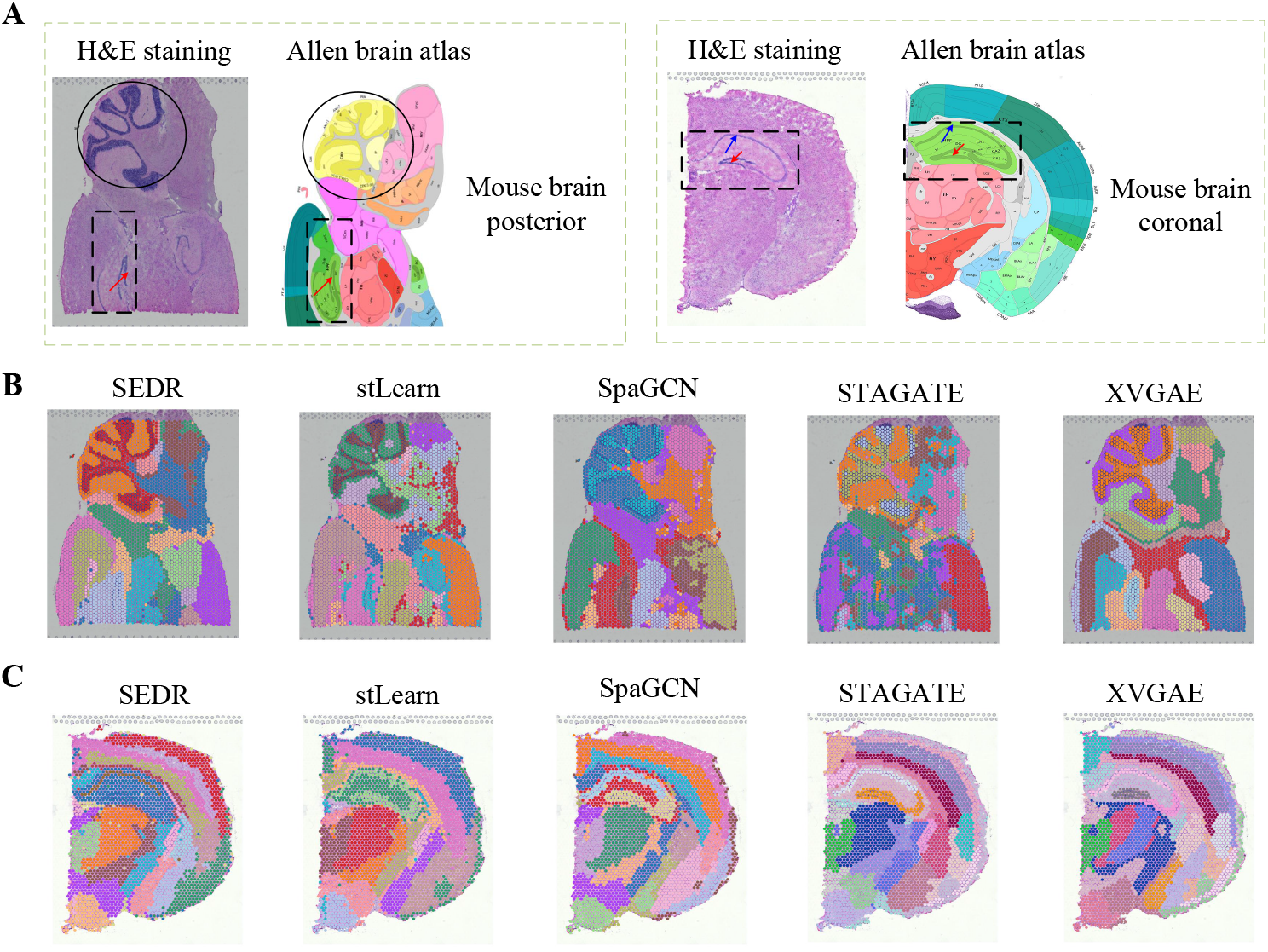
The XVGAE identifies better spatial domains for mouse brain dataset. (A) H&E staining of posterior and coronal regions of mouse brain tissue, and corresponding anatomical Allen Mouse Brain Atlas, where the circle denotes the cerebellar cortex region (CBX), the box denotes the hippocampus formation (HPF), the red arrows and the blue arrow denote the denate gyrus and the hippocampus proper in hippocampus formation, respectively. (B) and (C) are the visualization of clustering results for the Sagittal-Posterior and coronal slices of mouse brain, respectively, by XVGAE and other methods.

By comparing the spatial domains identified by these methods and the anatomical reference annotated by the Allen Mouse Brain Atlas [23], we found that in the posterior slice of the mouse brain, the SEDR, stLearn, and XVGAE methods could always accurately identify the cerebellar cortex region regardless of the chosen cluster resolution. Furthermore, the SEDR and XVGAE successfully identify the dentate gyrus region as the cluster number increases, shown in Fig. S6, indicating that they could effectively identify the hippocampal outline. We noticed that stLearn and SpaGCN methods could always identify the dentate gyrus region, while the other regions are not clear.

In coronal sections of the mouse brain, as shown in Fig. S7, we can observe the effects of different clustering resolutions compared with the Allen Mouse Brain Atlas. At lower clustering resolutions, all methods effectively distinguish various rough brain regions. However, at higher resolutions, such as in the cerebral cortex (CTX) and hippocampus formation (HPF) regions, the STAGATE and XVGAE methods excel in distinguishing different layers and accurately identifying the hippocampus itself. In contrast, SEDR, stLearn, and SpaGCN struggle to clearly delineate these regions, resulting in varying degrees of category ambiguity. As clustering resolution increases, stLearn, SpaGCN, STAGATE, and XVGAE progressively identify the dentate gyrus (DG) region, while SEDR shows limited recognition capability in this specific area. Additionally, in Fig. 3C, the XVGAE method clusters the hippocampus more clearly, distinguishing the DG and CA regions into separate classes. In comparison, STAGATE merges these regions into one class, demonstrating that our XVGAE method achieves superior clustering results.

We further evaluated the clustering results of different methods on the two datasets. Fig. S4 shows the silhouette coefficients of various methods for 10-25 clusters on both datasets. The silhouette coefficient value of XVGAE is significantly higher than that of other methods, quantitatively demonstrating that our method can delineate these regions more clearly. Additionally, we visualized the known marker genes of the two datasets in Fig. S5. For the MBP data, XVGAE effectively identifies the cerebellar cortex region and the dentate gyrus structure of the mouse brain. For the MBC data, XVGAE accurately captures the structure of the hippocampus region. These findings align with several specific marker genes presented in Fig. S5.

Overall, the results obtained by XVGAE on both the mouse brain posterior and mouse brain coronal are highly consistent with the annotations from the Allen Mouse Brain Atlas.

### 2.4 XVGAE accurately identifies the laminar organization of the mouse olfactory bulb tissue section profiled by Stereo-seq

We employed the XVGAE on a mouse olfactory bulb section profiled by Stereo-seq. As shown in Fig. 4A, [24] delineated the laminar architecture of the coronal mouse olfactory bulb section based on the DAPI-stained image. In Fig. 4B, we presented spatial domains identified by our XVGAE and other methods on this dataset. It is evident that XVGAE can distinctly discern various laminar regions and finely delineate boundaries, showing strong concordance with the annotation of the coronal mouse olfactory bulb, as provided by the Allen Reference Atlas [24]. On the contrary, other methods encounter confusion in specific areas and show a higher number of noisy points. For example, SEDR, SpaGCN, and stLearn could not identify the RMS and GCL layers, and the delineation of external areas is also unclear. The STAGATE accurately identifies the RMS and GCL layers, but unfortunately, it also confuses the GCL with the IPL layer.

**Fig. 4.**
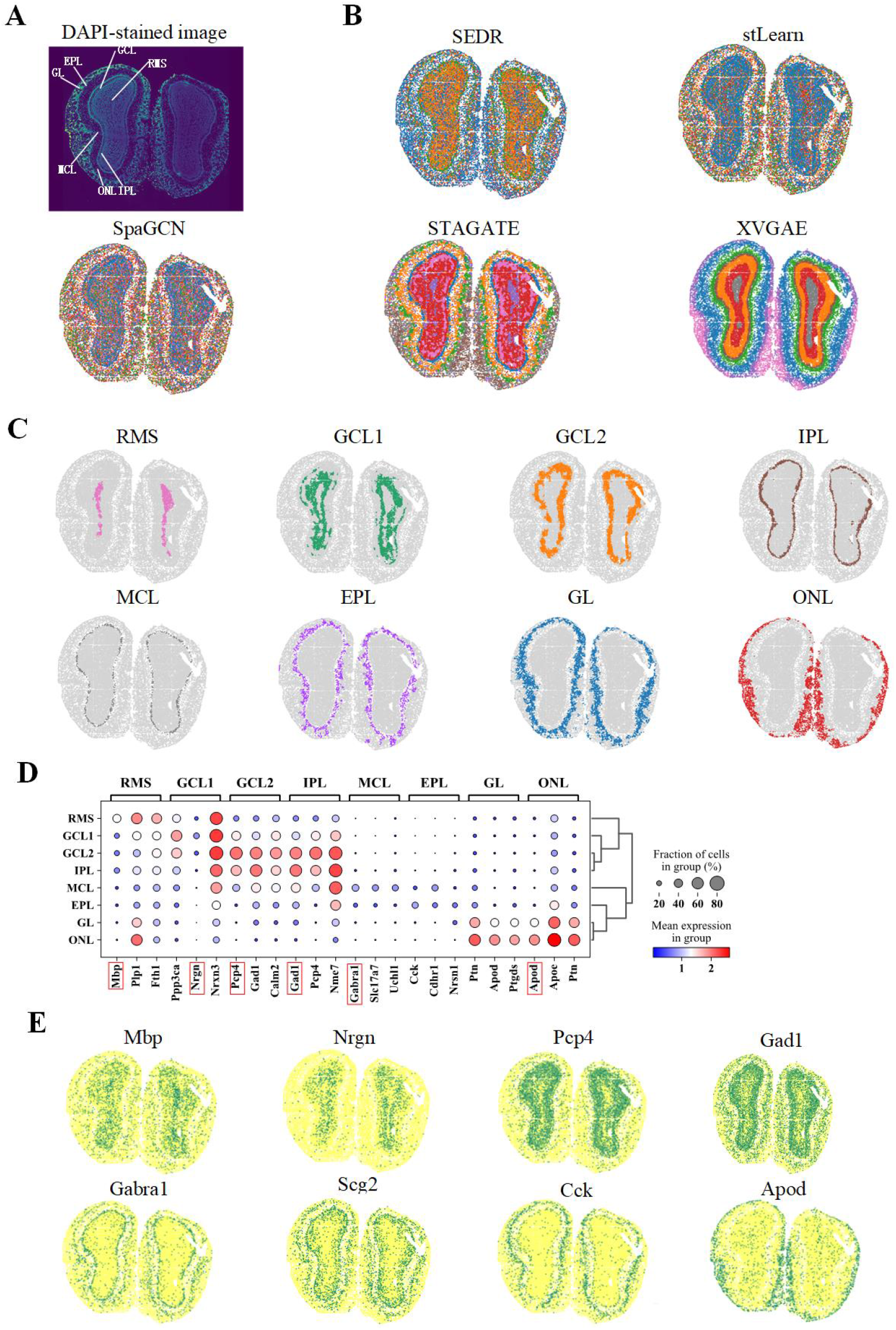
The laminar organization of the mouse olfactory bulb tissue section profiled by Stereo-seq was accurately identified by XVGAE. (A) The laminar organization of the mouse olfactory bulb is annotated in the DAPI-stained image produced by Stereo-seq. (B) Clustering results of XVGAE and compared methods on the Stereo-seq mouse olfactory bulb tissue section dataset, with the adjustment of Louvain resolution resulting in eight clusters. (C) Visualization of spatial domains identified by XVGAE layer by layer. (D) Differential genes between the spatial domains identified by XGVAE. (E) Visualization of the known marker genes for the spatial domains identified by XGVAE.

Figure 4C showcases the spatial domains identified by the XVGAE method, layer by layer. It is evident that XVGAE accurately delineates the boundaries of each layer, particularly the internal IPL and GCL layers, which the other methods fail to distinguish. Differential gene identification for the spatial domains identified by XVGAE is presented in Figure 4D. Some of these genes (encircled in red) are validated as known marker genes [25], and Fig. 4E further illustrates the spatial domains with their corresponding marker genes. We could see that the spatial domains are consistent with the performance of the known marker genes [25] illustrated in Fig. 4E. For example, XVGAE successfully identifies the compact MCL structure, characterized by the predominant expression of the mitral cell marker gene Gabra1 [26]. The stratified gene distribution exhibited by Nrgn, Pcp4 and Apod genes matches the GCL1, GCL2, and ONL layers in the clustering results, suggesting that the spatial domains revealed by XVGAE are robustly corroborated by established gene markers. Overall, this comprehensive analysis underscores the effectiveness of the XVGAE method in accurately identifying spatial domains within the tissue, as confirmed by the distribution of known marker genes.

### 2.5 XVGAE identifies the spatial domains within human breast cancer tumor tissue more accurately

To further assess the clustering performance and generalization capability of XVGAE in identifying cancer regions, we conducted experiments on a publicly available spatial transcriptomics dataset of breast cancer (invasive ductal carcinoma) [24] on the 10x Visium platform. The slices are manually annotated into 20 subregions and divided into four broad histological types based on pathological features [24]. We showed the histological images, annotations for spots types and pathological annotations in Fig. 5A.

**Fig. 5.**
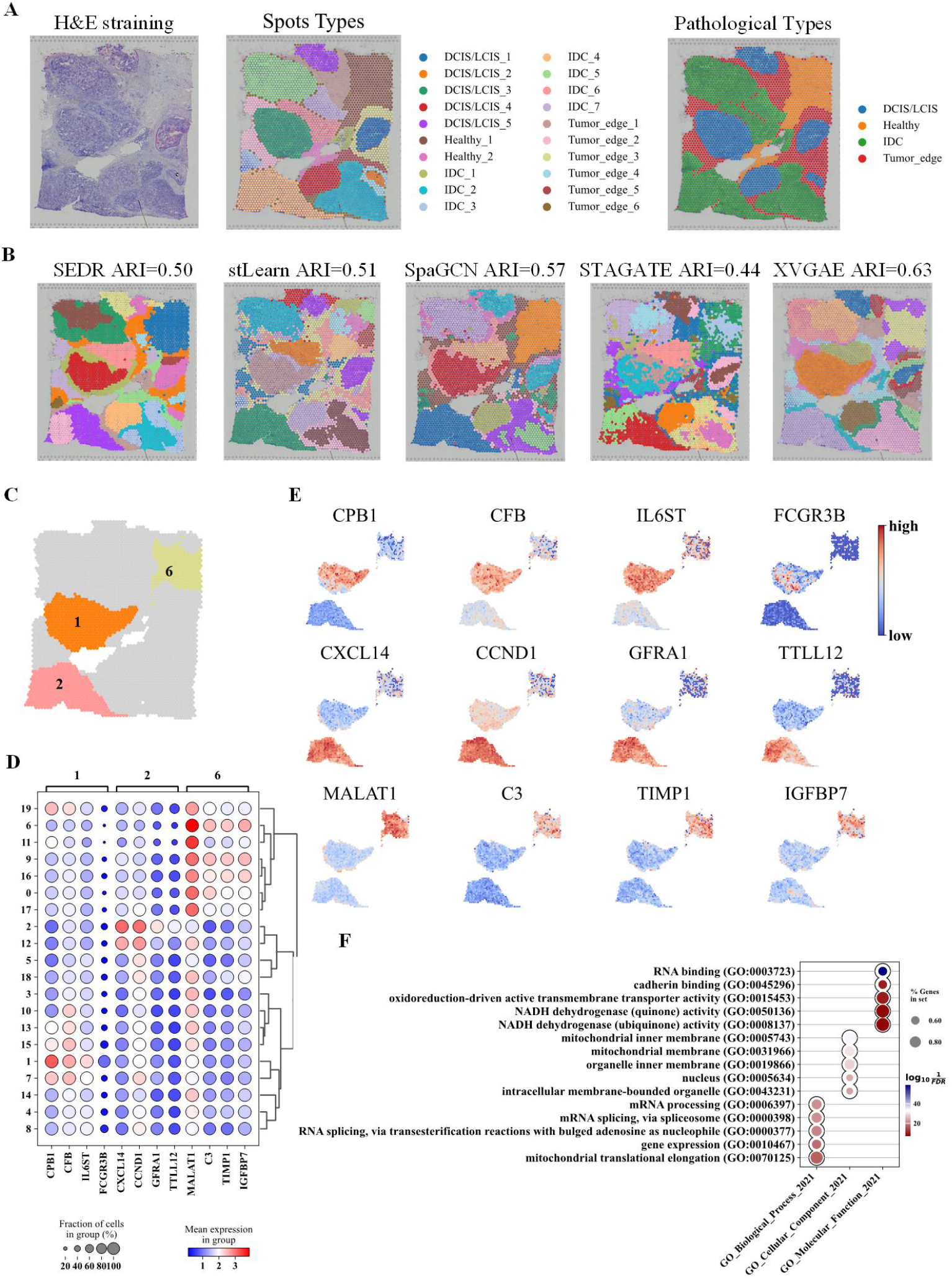
The XVGAE can dissect the spatial domains of human breast cancer tumor tissue at a more refined level. (A) The histological image, manual annotations for spots types and pathological types of the human breast cancer dataset are presented from left to right. For the pathological annotations, blue areas represent ductal carcinoma in situ and lobular carcinoma in situ (DCIS/LCIS), red areas represent invasive ductal carcinoma (IDC), yellow areas indicate healthy regions, and green areas signify the tumor edge. (B) Visualization of the clustering results for XVGAE and compared methods on the dataset. (C) Spatial locations for three sel1ec1ted domains 1, 2 and 6. (D) The differential gene dot plots of differential gene expression for Domains 1, 2 and 6, respectively. (E) Visualizations of differential gene expression for Domains 1, 2 and 6, respectively, from top to bottom row. (F) Gene ontology enrichment analysis of the differential genes between domains 2 and 6.

To quantify the clustering effectiveness of XVGAE on this dataset, we compared the identified spatial domains by different methods with the 20 manually annotated subregions. The identified spatial domains by XVGAE and other methods are shown in Fig. 5B. XVGAE obtains a higher adjusted random index (ARI) of 0.60 and delineates spatial domains with smoother and clearer boundaries, closely resembling the 20 manually annotated subregions, while comparison methods exhibit varying degrees of blurriness across spatial domains, often mixing spots of different types together. For instance, for the spatial locations corresponding to type IDC 4 in the lower left corner and type IDC 2 in the lower right corner of the ground truth, the compared methods tend to mark these areas as mixtures of multiple types, while XVGAE distinctly identifies the corresponding spatial domains with greater accuracy.

XVGAE also excels in spatial heterogeneity analysis. Tumor heterogeneity refers to the differences observed in various aspects such as growth rate, invasiveness, drug sensitivity, prognosis, etc., during the growth and division of tumor cells, which significantly influence the development of tumors. Therefore, investigating heterogeneity between tumors, and between tumors and healthy tissues is of great significance for gaining insights into the factors influencing tumor development and characteristics. To investigate the spatial heterogeneity of the breast cancer dataset, we selected three representative spatial domains for differential gene analysis as shown in Fig. 5C, namely Domain 1 (corresponding to DCIS/LCIS in the ground truth), Domain 2 (corresponding to IDC in the ground truth), and Domain 6 (corresponding to healthy tissue in the ground truth). The differential genes of the three domains are identified by the wilcoxon method, here we only reported the top four differentially expressed genes for the three domains in Fig. 5D.

Fig. 5E shows the expression values of the the top four DEGs for domains 1, 2 and 6, respectively. We could see significant upregulation of the CPB1 gene in domain 1, enabling a distinct separation of DCIS from other cancer regions. The CFB gene, involved in host defense and considered crucial in cellular damage and inflammatory processes, is also upregulated. As for Domain 2, the gene CXCL14 exhibits significant upregulation. The CXCL14 gene is involved in regulating various biological processes in the body, such as inflammatory immune responses, tumor-associated angiogenesis, activation of host-specific immunity against tumors, and autocrine regulation of tumor growth [27]. Studies have shown that the CXCL14 gene can enhance the invasiveness of breast cancer cells, thereby further promoting tumor development [28]. The CCND1 gene is also significantly upregulated in this domain. Mutations, amplifications, and overexpression of this gene can alter the cell cycle process and are commonly observed in various human cancers [29]. In Domain 6, the MALAT1 gene shows noticeable upregulation. Although once considered an oncogene, recent studies suggest that MALAT1 can inhibit breast cancer metastasis [30], now classifying it as a tumor-suppressor gene. The C3 gene, known for its regulatory role in inflammation and antimicrobial activity [31], is also identified. The TIMP1 gene, belonging to the TIMP gene family, encodes a protein considered a natural inhibitor of matrix metalloproteinases (MMPs) [32].

Furthermore, we reported results of gene ontology (GO) enrichment analysis based on the differential genes with adjusted *p−*values less then 0.05, for the domains 2 and 6 in Fig. 5F, and gene ontology enrichment analysis is conducted using GESApy (https://github.com/zqfang/GSEApy). In the subfigure, The rows represent enriched GO terms in three categories: BP (Biological process), CC (Cellular components) and MF (Molecular functions). The size of the circles represents the number of differential genes in the specific GO term, and the color represents the enrichment *p*-value.

From the enriched pathways in the subfigure, we can see that compared to the healthy region, the gene expression in the IDC region primarily manifests in the following five aspects: (i) Mitochondrial Function Alteration. We observed that several enriched GO terms involve mitochondrial functions, such as mitochondrial inner membrane (GO:0005743), mitochondrial membrane (GO:0031966), and mitochondrial translational elongation (GO:0070125). This indicates that the differential genes in invasive ductal carcinoma (IDC) may be associated with changes in mitochondrial function and structure. Mitochondria play a crucial role in cellular energy metabolism and apoptosis, and alterations in mitochondrial function could affect the energy supply and anti-apoptotic capabilities of cancer cells, where [33] has concluded through experiments that changes in cancer cell mitochondria may play a crucial role in the development of tumors. (ii) RNA Processing and Gene Expression Regulation.

Several enriched GO terms are related to RNA processing and gene expression regulation, such as mRNA processing (GO:0006397), mRNA splicing via spliceosome (GO:0000398), RNA splicing via transesterification reactions with bulged adenosine as nucleophile (GO:0000377), and gene expression (GO:0010467). These results suggest that differential genes in IDC may be involved in RNA splicing and processing, affecting gene expression regulation. Several studies [34–38] have shown that RNA processing and mRNA splicing play a key role in breast cancer. [39] found a significant induction of Tra2-*β*1 in invasive breast cancer, both on the RNA and protein levels. (iii) Changes in Oxidation-Reduction Reactions. The enrichment of GO terms such as oxidoreduction-driven active transmembrane transporter activity (GO:0015453) and NADH dehydrogenase (quinone) activity (GO:0050136) indicates changes in oxidation-reduction reactions in IDC. Oxidation-reduction reactions are crucial in cellular metabolism, and alterations in these reactions could lead to oxidative stress and metabolic reprogramming, thereby promoting cancer cell growth and invasion. For example, [40] pointed that aberration in mitochondrial complex I NADH dehydrogenase activity significantly enhance the invasiveness of human breast cancer cells, while therapeutic normalization of NAD+/NADH balance inhibits metastasis and blocks disease progression. (iv) Altered Cell Adhesion and Signaling. The enrichment of the GO term cadherin binding (GO:0045296) suggests that cell-cell adhesion and signaling may be altered in IDC. Cadherins play an important role in cell adhesion and maintaining tissue structure, and changes in their function could affect the migration and invasion capabilities of cancer cells. For example, [41] showed that EMT induction by E-cadherin loss promotes cancer cell invasiveness and that E-cadherin loss has broad transcriptional and functional effects in human breast epithelial cells. (v)Nucleus changes. The enrichment of the GO term nucleus (GO:0005634) indicates that nucleus function may be affected in IDC. In fact, as early as 1860, Beale observed changes in the size and shape of the nucleus in cancer cells [42]. Nuclear changes are a hallmark of many types of cancer, and a detailed understanding of the morphology and functional organization of the nucleus may help to understand the various changes in the cancer nucleus during the process of carcinogenesis [43].

In summary, by analyzing these enriched GO pathways, we can infer that differential genes in invasive ductal carcinoma may promote cancer cell growth, survival, and invasion through alterations in mitochondrial function, RNA processing and gene expression regulation, oxidation-reduction reactions, cell adhesion and signaling, and nuclear function. These inferences may provide clues for further studies on the molecular mechanisms of IDC and potential therapeutic targets.

## 3 Discussion

In this study, we have established a novel computational approach, XVGAE (cross-view graph autoencoders), designed for accurately identifying spatial domains in spatial transcriptomics data. XVGAE integrates spatial locations, gene expression profiles, and histological images, learning low-dimensional latent embeddings that enhance spatial domain identification. Traditional clustering methods rely heavily on gene expression data while neglecting spatial location information and histological images, often resulting in discontinuities and inaccuracies that do not align with the actual organizational structure of the tissue. More recent methods though improving clustering results often overlook the crucial histological information. To fully leverage histological information, XVGAE treats it as a distinct data modality and designs a cross-view graph convolutional networks for information integration. By constructing two graphs from spatial locations and histological images, and using the same gene expression levels as node features, cross-view GCNs allow XVGAE to learn specific representations for each view and propagate information between them, thus fully integrate multi-modal information for more accurate and biologically meaningful spatial domain identification.

The efficacy of XVGAE has been thoroughly validated using spatial transcriptomics data from multiple platforms. Comparative analysis with existing algorithms reveals that XVGAE exhibits strong capabilities in precisely identifying spatial domains, especially very thin domains. Despite the significant performance improvements, XVGAE encounters some limitations, for instance, model parameters needs to be readjusted when handling different tissue samples. Additionally, the method may pose challenges when processing extremely large datasets. Future research could focus on further optimizing XVGAE to enhance its adaptability and performance across diverse datasets. In addition, current computational methods in the field of image processing demonstrate robust performance. Processing gene expression and spatial location information as image views could potentially enhance the accuracy of spatial domain identification. Overall, integrating histological information and employing multi-view learning algorithms to improve the resolution and accuracy of spatial transcriptomics data are promising directions. These advances can improve the accuracy of deep learning-based reconstructions and may help provide some insights for applications in biological and medical research.

## 4 Conclusion

Accurately identifying spatial domains is crucial for understanding cellular function and tissue conditions in both physiological and pathological states. In this work, we developed novel cross-view graph autoencoder framework for spatial domain identification called XVGAE. XVGAE leverages cross-view autoencoders that integrate a histological graph and a spatial graph with shared nodes representations to learn more informative latent representations. It enhances the representations through self-optimized clustering. Through benchmark comparisons with five baseline methods on four real spatial transcriptomics datasets over different platforms, XVGAE shows better performance across various metrics. For datasets featuring samples with clear boundaries and complex tissues, XVGAE identifies spatial domains that are more consistent with known structures. Our experiments showed that XVGAE excels in improving the separation of small and thin spatial domains compared to other methods. This improvement allows for a better characterization of the spatial expression patterns of domain-specific genes, thereby enhancing our understanding of tumor heterogeneity and advancing personalized treatment approaches for cancer. In summary, regardless of tissue complexity, domain size, or the resolution of spatial transcriptomics technology, XVGAE exhibits excellent generalizability across datasets.

## 5 Methods

XVGAE takes the spatial transcriptomics dataset as input, specifically including three components: *X*_0_ ∈ *R*^*N*×*d*^, representing the gene expression matrix for *d* genes and *N* spots, *P* ∈ *R*^*N*×2^, representing the two-dimensional spatial coordinates of each spot, and the histological image *I*. The overview of the XVGAE is shown in Fig. 1.

### 5.1 Data preprocessing

The original gene expression data *X*_0_ is normalized using the scanpy [44] software package. We then select the top 3000 highly variable genes and conduct PCA for dimension reduction, resulting in a matrix *X* ∈ *R*^*N*×50^. For the histology image *I*, we obtain visual features *X*_*I*_ ∈ *R*^*N*×30^ using the SimCLR model [45], which maximizes agreement between different augmented views of the same spot image via a contrastive loss in the latent space.

### 5.2 Construction of the spatial graph and histological graph

The relationship among the spots could be captured from either the spatial locations or the histological image. We construct two graphs from the spatial locations and the histological images, respectively, for the spot nodes with the same gene expression profiles as node attributes. The adjacent matrices *A*_*P*_ and *A*_*I*_ for the two graphs are constructed by K-nearest neighbors (KNN) algorithm, based on Euclidean distances derived from the spatial locations *P* and preprocessed histological image features *X*_*I*_, respectively. Two attributed graphs, *G*_*P*_ = (*A*_*P*_, *X*) and *G*_*I*_ = (*A*_*I*_, *X*) for the spots are constructed, from the spatial view and the histological view, respectively.

### 5.3 Cross-view graph autoencoders

In this section, we propose a cross-view graph autoencoders framework to fully integrate the two attributed graphs 𝒢_*P*_ and 𝒢_*I*_ from the spatial view and histological view. The XVGAE consists of a cross-view encoder to capture the hidden representations for spots by fully integrating the two graphs, along with a decoder for reconstructing the gene expression matrix *X*.

For the cross-view encoder, two graph convolutional networks (GCNs) are first taken to extract the view-specific features for spatial graph 𝒢_*P*_ and histological graph 𝒢_*I*_ by

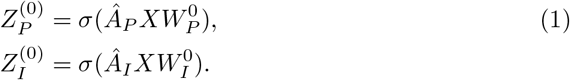

where *σ* is the activation function set as the rectified linear unit(ReLU) [46], and 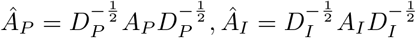are the normalized adjacency matrice, *D*_*P*_ and *D*_*I*_ are diagonal matrices with row sums of *A*_*P*_ and *A*_*I*_, respectively, and 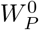 and 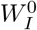 are model parameters that need to be learned.

To fully integrate the view-specific information from both views, the cross-view encoder further captures and conducts two cross-view GCNs on the adjacency matrix from one view with view-specific representations from the other view. Specifically, to update the view-specific representations 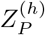 and 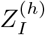 for *h*-th layer, one GCN is applied on the adjacency matrix *Â*_*P*_ from the spatial view and the view-specific representation 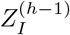 from the histological view, and the other GCN is on the adjacency matrix *A*_*I*_ from the histological view and the view-specific representation 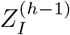 from the spatial view, as follows,

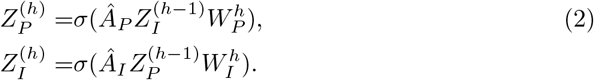

The cross-view GCNs enable the mutual propagation of the view-specific information between the spatial view and the histological view. After *L* layers, the average of 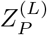 and 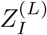 is taken as the fused representations *Z*_*G*_ for the two graphs.

Besides the cross-view graph encoder that utilizes the graph information, the gene expression features *Z*_*X*_ are also directly extracted from the gene expression matrix *X* by an encoder with a three-layer fully connected neural network. Subsequently, the combination of *Z*_*G*_ and *Z*_*X*_ yields the final representation:

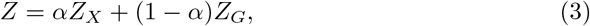

where *α* is a learnable coefficient, which is chosen based on the properties of the respective dataset. In this study, *α* is initialized to 0.5 and then automatically adjusted using a gradient descent approach.

Finally, the decoder is a three-layer fully connected neural network that recovers the gene expression matrix 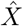 by the fused representations *Z*.

### 5.4 Loss function

The loss function for XVGAE consists of a reconstruction loss *L*_*REC*_, which accounts for the gene expression reconstruction, and a clustering loss *L*_*KL*_, which accounts for the clustering structure in the fused representations *Z*.

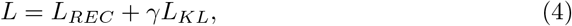

where *γ* is a hyperparameter that balances the importance of reconstruction and clustering terms and is set to 0.5 by default.

Specifically, the reconstruction loss of normalized expressions can be written as follows:

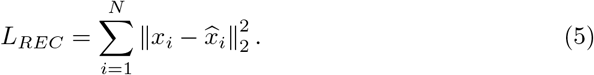

For the clustering loss, we employed an unsupervised deep-embedded clustering method to iteratively group spots into distinct clusters from learned latent representations *Z*. The K-means algorithm [47] is used to initialize the cluster centers based on *Z*. Let *C* be the number of initial clusters. In the first step, a soft assignment *q*_*iu*_, *u* = 1, 2, *·, C* is computed using the Student’s t-distribution, which serves to evaluate the similarity between the spot embedding *z*_*i*_ and cluster center embedding *µ*_*u*_:

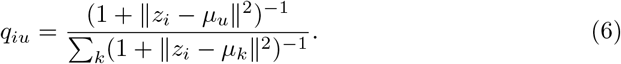

The clusters are then iteratively refined by learning from the high-confidence assignments to an auxiliary target distribution *p*, based on the following formula:

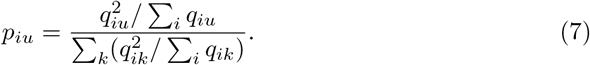

By employing the soft assignment *q*_*iu*_ and auxiliary target distribution *p*_*iu*_, the clustering loss function is defined using the KL divergence:

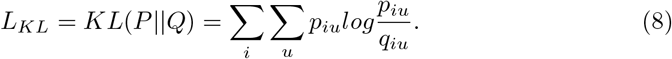

## 6 Competing interests

No competing interest is declared.

## 7 Data availability

The code and detailed tutorial for XVGAE can be accessed through the following link: https://github.com/LiminLi-xjtu/XVGAE. All utilized datasets are publicly available for easy download:

(1) DLPFC: The dataset comprises 12 slices, each containing varying spot quantities ranging from 3460 to 4789. The download link is as follows:http://spatial.libd.org/spatialLIBD
(2) Mouse brain: The dataset comprises mouse brain data, encompassing two sections: the anterior slice and the posterior slice. The download link is provided below:https://support.10xgenomics.com/spatial-gene-expression/datasets
(3) Human breast cancer: The breast cancer dataset contains 3798 spots and 4 main histological types. Human breast cancer tissue sections datasets are available at https://support.10xgenomics.com/spatial-gene-expression/datasets.
(4) Mouse olfactory bulb: Mouse olfactory bulb tissue datasets are available at https://github.com/JinmiaoChenLab/SEDR.

## 8 Acknowledgments

This work was funded by the National Natural Science Foundation of China project under Grant No. 12222115.

## Notes

### Competing Interest Statement

The authors have declared no competing interest.

