## Supplementary Materials for "Cross-view graph neural networks for spatial domain identification by integrating gene expression, spatial locations with histological images"

#### 1 Supplementary note for Clustering metrics

**Adjusted rank index(ARI)**

For datasets with ground truth labels, we use adjusted rank index (ARI) [1] to compare the performance of different clustering algorithms. The ARI score is defined as

$$ARI = \frac{\sum_i \binom{n_{ij}}{2} - \left[\sum_i \binom{p_i}{2} \sum_j \binom{g_j}{2}\right] / \binom{N}{2}}{\frac{1}{2}\left[\sum_i \binom{p_i}{2} + \sum_j \binom{g_j}{2}\right] - \left[\sum_i \binom{p_i}{2} \sum_j \binom{g_j}{2}\right] / \binom{N}{2}}, \tag{1}$$

where $N$ is the total number of spots, $p_i$ is the number of spots assigned to $i$-th calculated cluster, $g_j$ is the number of spots belonging to the $j$-th ground truth cluster, and $n_{ij}$ is the number of overlapping spots between the $i$-th calculated cluster and the $j$-th ground truth cluster. A higher ARI score implies better clustering performance.

**Silhouette coefficient**

If there are no available spatial domain annotations, the commonly used clustering metric is the silhouette coefficient (SC score). The formula for calculating the SC score is as follows:

$$\mathrm{S(i)} = \frac{\mathrm{b(i) - a(i)}}{\max\{\mathrm{a(i), b(i)}\}}, \tag{2}$$

where a(i) is defined as $\mathrm{a(i)} = \frac{1}{\mathrm{n-1}} \sum_{\mathrm{j=1}}^{\mathrm{n}} \mathrm{distance(i, j)}$, which calculates the average distance between each sample $i$ and all other samples within the same cluster, b(i) calculates the average distance between each sample $i$ and all samples in the nearest neighboring cluster. The SC score ranges from -1 to 1 and a higher value suggests that the clusters are dense and well-separated.

**Algorithm 1** Algorithm XVGAE

1: Input spatial transcriptomics data with gene expression $X_0$ and spatial location $P$, and histological image $I$.
2: Data preprocessing: for gene expression data $X_0$, normalize, select highly variable genes, and use PCA method to obtain the low-dimensional embedding $X \in R^{N\times 50}$; for histological image $I$, obtain visual features $X_I \in R^{N\times 30}$ using the SimCLR model [2].
3: Constructing attributed graphs: use K-nearest neighbors (KNN) algorithm to obtain adjacent matrices $A_P$ and $A_I$ for spatial graph and histological graph based on $P$ and $X_I$, respectively, the corresponding attributed graphs are: $\mathcal{G}_P = (A_P, X)$ and $\mathcal{G}_I = (A_I, X)$.
4: Use two graph convolutional networks (GCNs) to extract the view-specific features $Z^{(}_{P}0)$ and $Z^{(}_{I}0)$ by eq (1).
5: **for** $i = 1$ to $L$ **do**
6: Integrate the view-specific information from both views by the cross-view encoder in eq (2).
7: Calculate the loss fuction in eq (4) and update parameters.
8: **end for**
9: Average $Z_P^{(L)}$ and $Z_I^{(L)}$ as the fused representations $Z_G$ for the two graphs.
10: Extract the gene expression features $Z_X$ from $X$, and combine $Z_G$ and $Z_X$ as the final representation by eq (3).
11: Output: Perform clustering for the final representation $Z$ by k-means algorithm.

### 2 Supplementary Notes for algorithm XVGAE

### 3 Supplementary Notes for comparison methods

We compared XVGAE with six recently developed spatial-aware clustering approaches including SEDR [3], stLearn [4], SpaGCN [5], BayesSpace [6] and STAGATE [7] The parameter settings of these methods are as follows:
**(1) SEDR:** We ran SEDR for all experiments with its recommended parameters in the project on github (run_SEDR_DLPFC_all_data.py).

- Parameters setting: We ran these methods with the recommended parameters : k is set as 10, the epoch is set as 200, using_dec_loss is set as true.

**(2) stLearn:** We ran stLearn for all experiments with its recommended parameters in the project on github (DLPFC_stLearn.py).

- Parameters setting: We do the preprocessing for gene count table by running st.pp.filter_genes(), st.pp.normalize_total() and st.pp.log1p(),and we run st.em.run_pca() for gene expression data.

**(3) SpaGCN:** We ran SpaGCN for all experiments with its recommended parameters in the project on github (DLPFC_SpaGCN.py)

- Parameters setting: We ran these methods with the recommended parameters: s=1, b=49, p=0.5, max_epochs=200, and we run spg.prefilter_genes() and spg.prefilter_specialgenes() to avoid all genes are zeros. We run sc.pp.normalize_per_cell() and sc.pp.log1p() to normalize and take log for UMI.

**(4) BayesSpace:** We ran BayesSpace for all experiments with its recommended parameters in the project on github (DLPFC_BayesSpace.R)

- Parameters setting: We ran these methods with the recommended parameters: PCA_number is set as 15, top genes is set as 2,000, seed is set as 104.

**(5) STAGATE:** We ran BayesSpace for all experiments with its recommended parameters in their online tutorial (https://stagate.readthedocs.io/en/latest/index.html).

- Parameters setting: We run sc.pp.highly_variable_genes(), sc.pp.normalize_total() and sc.pp.log1p() to normalize the gene data, the n_top_genes is set as 3000, target_sum is set as 1e4.

**(6) XVGAE:** We give parameter settings of XVGAE on all test datasets.

- Human dorsolateral prefrontal cortex (DLPFC): n_componentsis set as 50, n_top_genes is set as 3000, target_sum is set as 1e4, epoch_pre is set as 200, epoch is set as 400, rad_cutoff=150, k_cutoff=8, n_clusters determined based on ground truth, hidden_size is set as 128, embedding_size is set as 20, gamma_value is set as 10.
- Human breast cancer (BRCA): all parameter settings are same as DLPFC datasets, as above, but rad_cutoff=500, k_cutoff=8.
- Mouse brain tissue: all parameter settings are same as DLPFC datasets, as above, but n_clusters is set as 10 to 25, rad_cutoff=250, k_cutoff=8.
- Mouse olfactory bulb tissue: all parameter settings are same as DLPFC datasets, as above, but we use louvain to complete clustering.

### 4 Supplementary figures

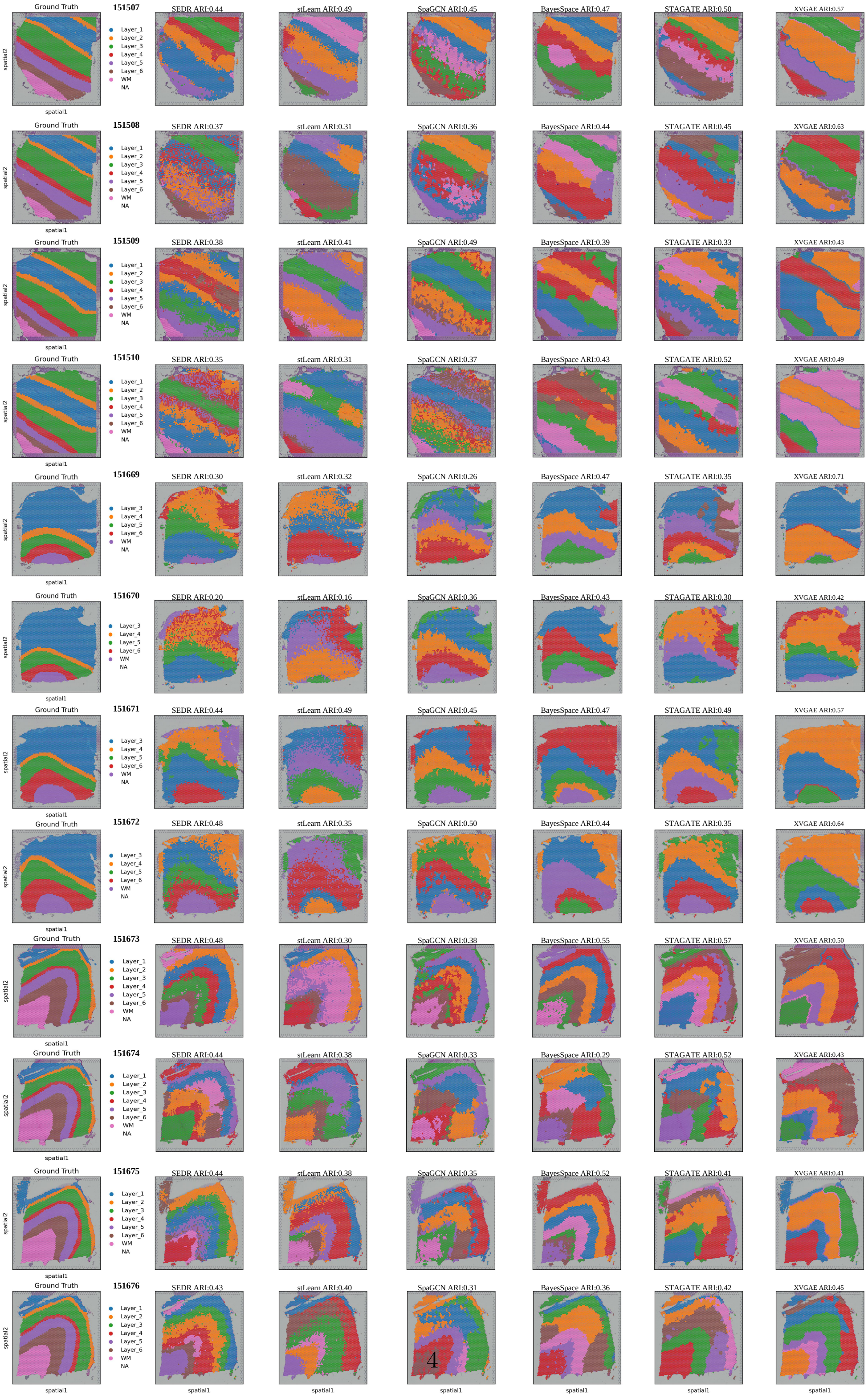




**Fig. S1** The spatial domains of the ground truth by manual annotations and the spatial domains that are identified by SEDR, stLearn, SpaGCN, BayesSpace, STAGATE and XVGAE for all 12 sections of the DLPFC dataset.

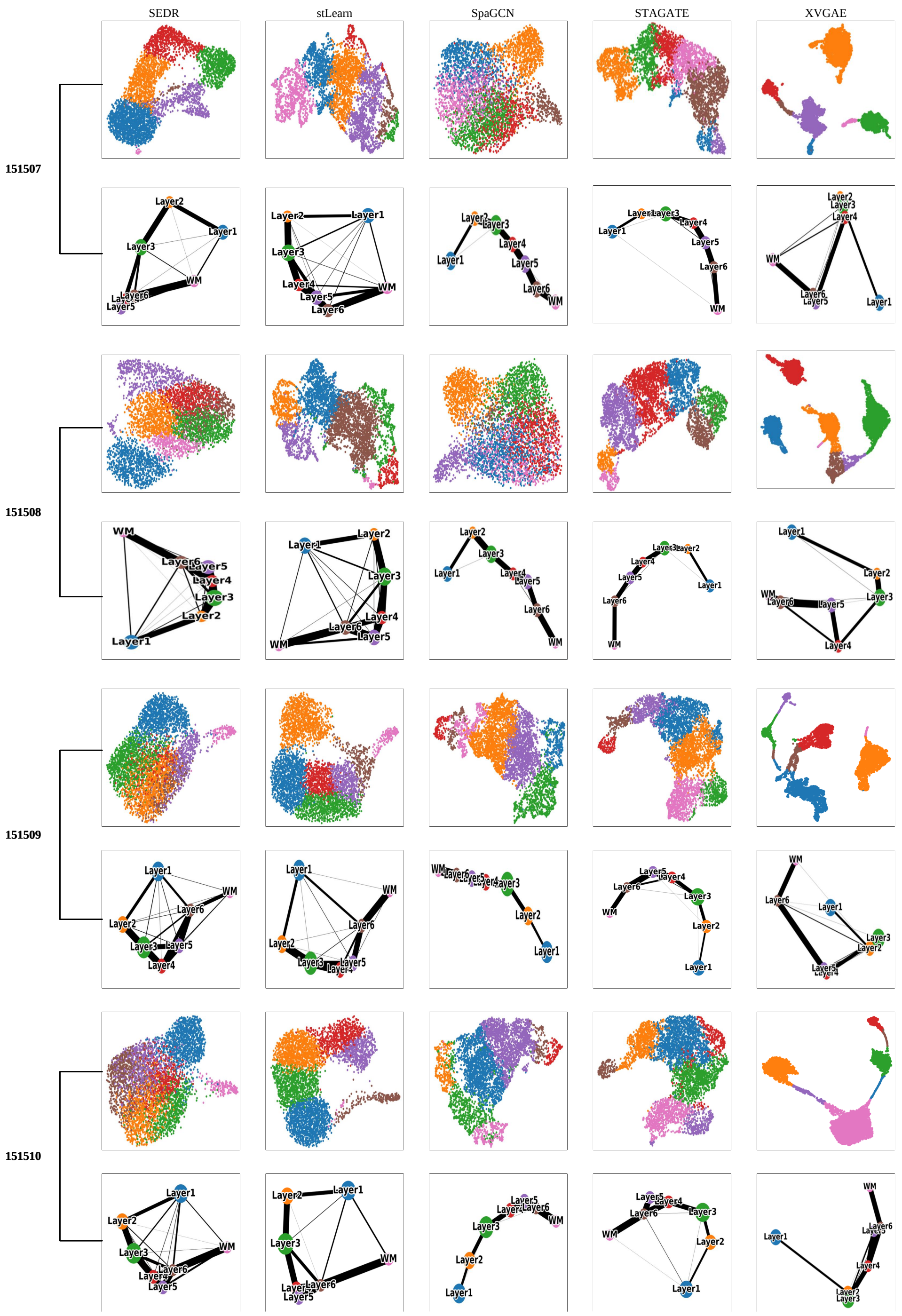
SEDR
stLearn
SpaGCN
STAGATE
XVGAE
151507
151508
151509
151510

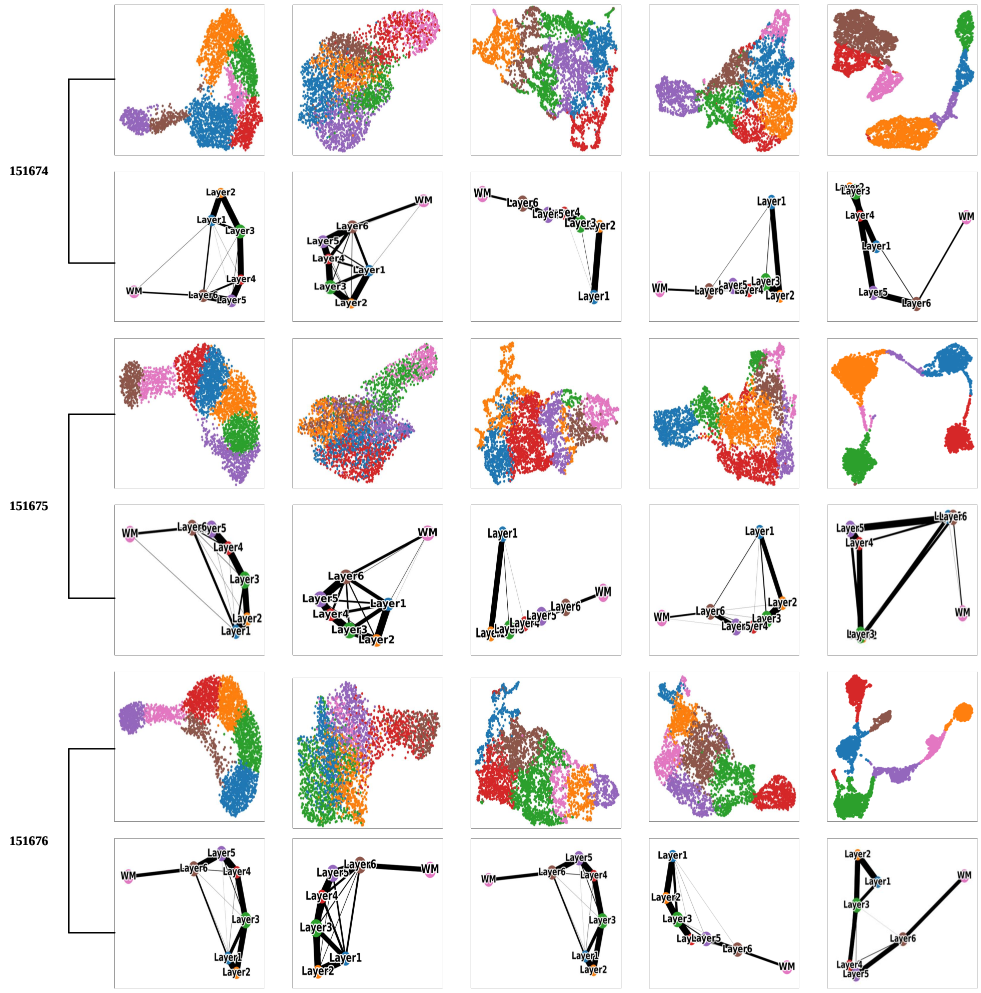


**Fig. S2** UMAP and PAGA visualization generated by the SEDR, stLearn, SpaGCN, STAGATE and XVGAE embeddings, respectively, for 12 sections of the DLPFC dataset.

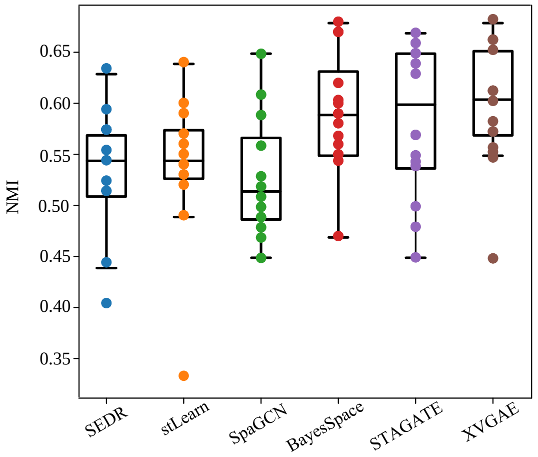


**Fig. S3** NMI boxplots of the performance of XVGAE and other algorithms for all 12 sections of the DLPFC dataset. Within the boxplot, the central line, box boundaries, and whiskers represent the median, upper and lower quantiles, and 1.5 times the interquantile range, respectively.

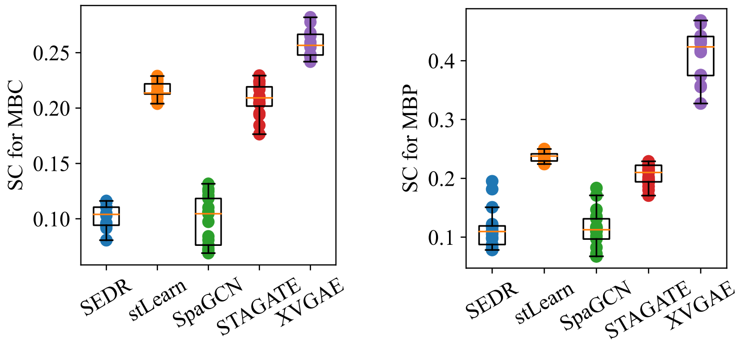


**Fig. S4** Histograms of Silhouette Coefficients (SC) scores by different methods for Mouse brain posterior (left) and Mouse brain coronal (right) datasets, respectively.

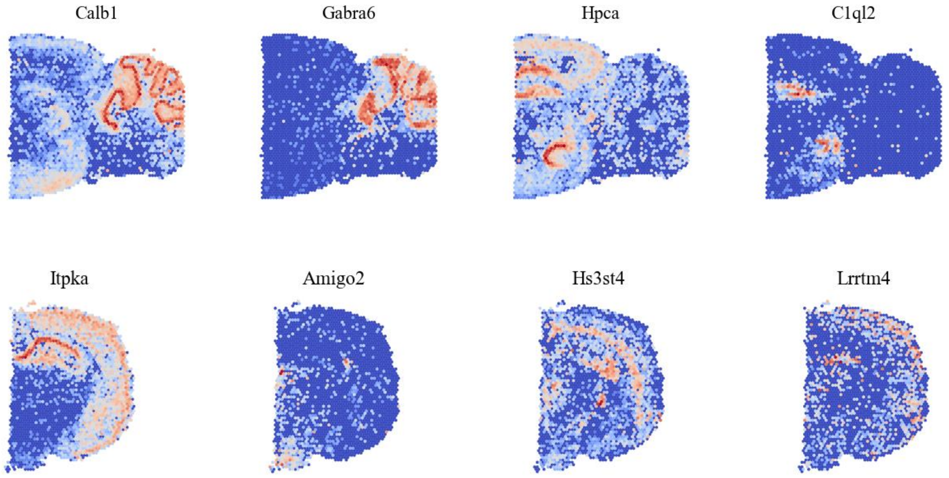


**Fig. S5** Expression levels for several marker genes for Mouse brain posterior (first row) and Mouse brain coronal (bottom row) datasets, respectively.

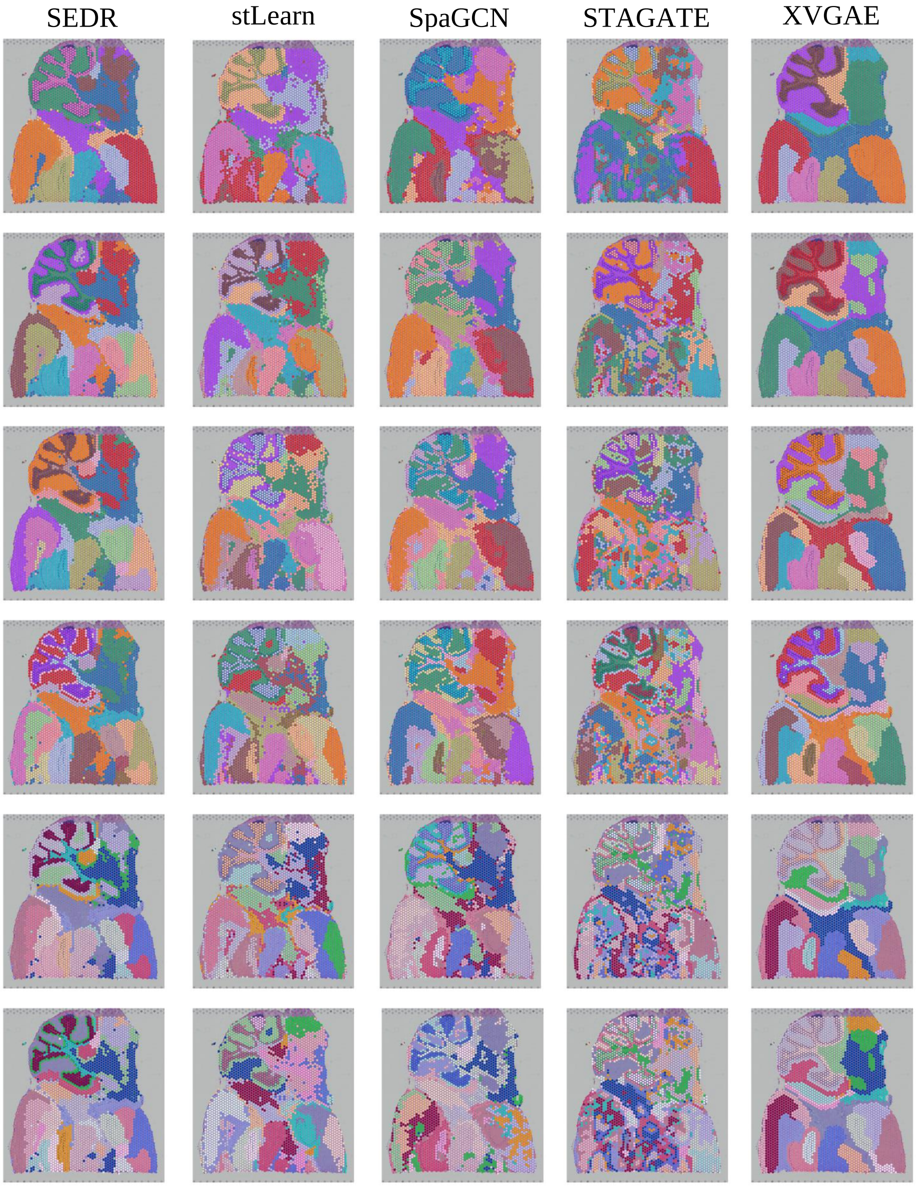


**Fig. S6** Spatial domains identified by SEDR, stLearn, SpaGCN, BayesSpace, STAGATE and XVGAE on the Mouse brain posterior data with different clustering numbers 11, 15, 17, 20, 22 and 24 from top to bottom.

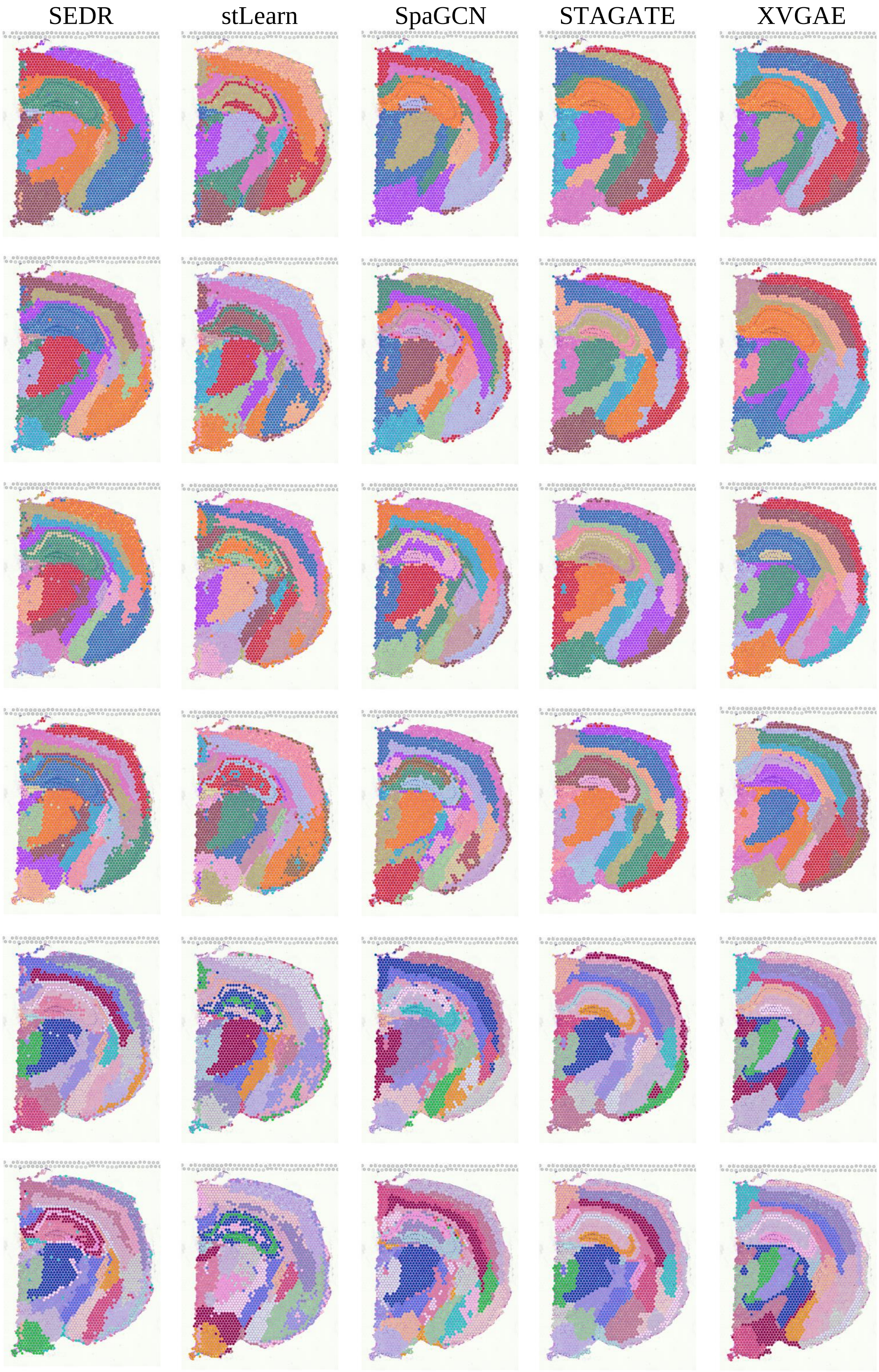




**Fig. S7** Spatial domains identified by SEDR, stLearn, SpaGCN, BayesSpace, STAGATE and XVGAE on the Mouse brain coronal data with clustering numbers 11, 15, 17, 20, 22 and 24 from top to bottom.
